# A 2-Compartment Bone Tumor Model for Testing the Efficacy of Cancer Drugs

**DOI:** 10.1101/829879

**Authors:** Aylin Komez, Arda Buyuksungur, Ezgi Antmen, Wojciech Swieszkowski, Nesrin Hasirci, Vasif Hasirci

## Abstract

We produced a three dimensional (3D) bone tumor model (BTM) to study the interactions between healthy and tumor cells in a tumor tissue microenvironment, migration of the tumor cells and the efficacy of an anticancer drug, Doxorubicin, for personalized medicine applications. The model consisted of two compartments: (a) a healthy bone tissue mimic, poly(lactic acid-co-glycolic acid) (PLGA)/beta-tricalcium phosphate (β-TCP) sponge that was seeded with human fetal osteoblastic cells (hFOB) and human umbilical vein endothelial cells (HUVECs), and (b) a tumor mimic, a lyophilized collagen sponge that was seeded with human osteosarcoma cells (Saos-2). The tumor component was introduced to a central cavity created in the healthy bone mimic and together they constituted the total 3D model (3D-BTM). The scaffolds were characterized by determining their mechanical properties, studying their topography and stability with compression tests, microCT, SEM, confocal microscopy and gravimetry. Porosities of the sponges were determined from µCT data as 96.7% and 86% for PLGA/TCP and collagen sponges, respectively. The average diameters of the pores were measured by using ImageJ (NIH, USA) as 199±52 µm for PLGA/TCP and 50-150 μm for collagen scaffolds. Young’ modulus of the PLGA/TCP and collagen sponges were determined as 4.76MPa and 140kPa, respectively. Cells seeded on the two sponges were studied independently and together on the BTM. Cell proliferation, morphology, calcium phosphate forming capacity and ALP production were studied on both healthy bone and tumor mimics. All types of cells showed cellular extensions and spread on and in the scaffolds indicating good cell-material interactions. Angiogenic developments in BTM were studied along with migration of cells between the components with immunocytochemistry, SEM, microCT, qRT-PCR and agarose gel electrophoresis. Confocal microscopy showed that a direct contact was established between the cells present in different parts of the BTM; and the HUVEC cells within the healthy bone mimic were observed to migrate into the tumor mimic. This was confirmed by the increase in the levels of angiogenic factors VEGF, bFGF, and IL-8 in the tumor component. The IC_50_ of Doxorubicin on Saos-2 cells was determined as 0.1876 µg.mL^−1^. Doxorubicin was administered to the BTM at 2.7 µg.mL^−1^ concentration and after allowing one day for interaction, the cell number was determined with Alamar Blue cell viability test as 7-fold lesser compared to 24 h earlier. Apoptosis of the osteosarcoma cells was measured by caspase-3 enzyme activity assay. These results demonstrate the suitability of the 3D BTM model for use in the investigation of activities and migrations of cells in a tumor tissue. These will be very useful in studying metastatic capabilities of cells in addition to personalized drug treatments.

## 1. Introduction

Cancer is a challenging health issue since the clinical success rate is very low. Presence of various cancer types, occurrence at different sites in the body and metastasis to other tissue or organs make the treatment difficult. The tumor may be benign or malignant. Cells of benign cancers do not spread to other tissues; meanwhile, malignant cells can migrate and metastasize. Most of the deaths are due to metastases occurring by migration of cancer cells via vascular or lymphatic system. Vascular or lymphatic system invasion happens when cancer cells break into blood vessels or lymph channels.

In case of bone, benign tumors do not metastasize but damage the bone structure and lower the mechanical strength.(1) One of the most prevalent primary malignant bone tumor is osteosarcoma (OS) and is primarily observed in children and young adults.(2) The survival rate in 5 years is 75% for bone cancers that do not spread from the primary site (3), but for the metastatic case general survival rate is 40% and in poor responders to chemotherapy the survival rate is lower, 20%.(3,4). In order to develop an effective therapy, a model system that closely mimics the tumor of the patient would be very valuable, especially in selecting drugs and dose regimens for that specific patient, in other words, for personalized therapy.(5)

We also need tissue models which mimic the complex organization of the natural tissue the extracellular matrix (ECM) and the microenvironment of the cancer cells in the tumor area in order to understand the underlying mechanisms in cancer occurrence and to develop rational approaches for diagnosis and therapy. In the last five decades, mainly two-dimensional (2D) monolayer tissue cultures were used to simulate the cancer cell microenvironment and study cancer.(6) These 2D systems were simple, easy to construct and convenient to study cell behavior (viability, topography, expressed proteins, etc.). However, these 2D systems have higher oxygen, glucose and nutrient concentrations, and therefore, can poorly mimic the natural tumor microenvironments,(7). It is also reported that the gene expression, topology, and biochemistry of the cells change in 2D cultures.(8) In addition, in 2D monolayer cultures the drugs tested show their toxic effects on the cells far more rapidly than in the natural 3D environment of the cancer tissue.(9) In the absence of interactions with the surrounding stroma as it is in the 2D environment, cells growing adherently lose their polarity which also alters several cell responses, such as apoptosis. All these reasons mentioned above led the researchers to make incorrect conclusions as in the case in most biologically targeted therapies which perform well in the lab but not in the clinics.(10) As a result, 3D organotypic models which better mimic the properties of the tumor tissue have become the preferred model types. Some researchers developed 3D models involving spheroids formed by spontaneous aggregation of the cells on agarose,(11) on poly(2-hydroxyethylmethacrylate) (pHEMA) coated plates,(12) or on ultralow attachment culture plates.(13) There are also matrix-based 3D tumor models. Ewing’s sarcoma cells cultured on electrospun poly(ε-caprolactone) (PCL) mats was shown to mimic human tumors in terms of growth and expression levels of therapeutically targeted genes present in crucial signaling pathways.(14) Models constructed using osteosarcoma cells grown on silk fibroin scaffolds were capable of expressing specific surface markers (HIF-1a and bFGF) at levels close to that of xenograft tumors.(15) U2OS cell line (osteosarcoma) cultured in 3D collagen matrices activated ‘Phosphatide Inositol-3 Kinase (PI3K) pathway’ which is a critical intracellular signaling cascade influencing cell growth, migration, protein expression, and survival.(16) Even though these models attempted to mimic the 3D tumors, the healthy bone microenvironment that has critical importance in the tumor initiation and progression were not taken into consideration. In one study, the importance of tumor microenvironment was shown in a model of human bone cancer (Ewing’s sarcoma, ES) where tumor cell spheroids were cultured in a bone tissue environment consisting of human mesenchymal stem cells in decellularized bone matrix. Their results demonstrated that cancer cells re-expressed focal adhesion and cancer-related genes that are normally highly expressed in tumors but they lost this property in monolayer cultures and gained angiogenic capacity that favor tumor initiation and progression.(17) They have taken their 3D construct one step further by developing a model containing both osteoclasts and osteoblasts within 3D mineralized bone matrix to mimic the *in vivo* bone osteolysis associated with the Ewing’s sarcoma to understand the mechanisms underlying tumor progression and also for the development of new therapeutics. Culturing of ES cell spheroids in the bone microenvironment led to a decrease in bone density, connectivity, and matrix deposition. Additionally, a therapeutic reagent which have demonstrated efficacy in ES treatment, prevented bone resorption mediated by osteoclasts in the tumor model.(18) This model appeared suitable to understand the biology and genetic origin of tumor and underlying reasons of tumor progression. However, choice of the appropriate drug for personalized therapy and discovery of new agents in the treatment of bone cancer require reproducible models unlike the decellularized bone matrix-based models that are not obtained in the same composition for each test. It is important to develop biomimetic tumor models that involve the primary components required for the precise mimicking of *in vivo* conditions that are not exceedingly complex and, therefore will not lead to complex data analysis.(19)

The main hypothesis of this study was that a bone tumor model could be constructed with a rigid bone-like exterior carrying healthy cells and a softer tumor tissue-like core carrying cancerous cells. In order to reflect the complex structure of bone, three biomaterials were selected; poly(lactic acid-co-glycolic acid) (PLGA) and β-tricalcium phosphate (β-TCP) as the organic and inorganic components of the healthy exterior region, and collagen as the tumorous core of the model. To be able to closely mimic the complex structural and mechanical properties of bone, these biomaterials were used since they represent both the bone mineral and the softer tumor tissue better than any new polymer family.

One of our main aims was to mimic the interaction between healthy and tumor bone cells as in their natural microenvironment. Bone tumor model was constructed with well established biomaterials and with a naturally mimicking design to study the migration of healthy bone cells and tumor cells and interactions between them.

The 3D bone tumor model (BTM) developed in this study has two tissue engineered components; one mimics the tumor tissue and the other, the healthy bone environment surrounding the tumor. A cylindrical collagen sponge was seeded with human osteosarcoma cells (Saos-2) and used as the tumor tissue mimic. PLGA sponge containing β-TCP (PLGA/TCP) was seeded with both human fetal osteoblastic cells (hFOB) and human umbilical vein endothelial cells (HUVECs) and constituted the healthy bone mimic. The collagen-based tumor mimic was inserted into a cavity at the center of the cylindrical shaped healthy bone mimic to form the complete bone tumor model (BTM). The interaction, communication, and migration of the cells in and between the two compartments of the BTM model were investigated. Then, the model was used to study the efficacy of an anticancer drug, Doxorubicin, in treating cancer in the tumor mimic. One of the future applications of this model would be to employ it for personalized cancer therapy, especially in the determination of the effective drug types and doses using cancerous cells of the patient.

## 2. Materials and Methods

### 2.1. Preparation of Sponges and 3D Bone Tumor Model

#### 2.1.1. Preparation of PLGA/TCP Sponges (scaffold of the healthy bone mimic)

Porous PLGA/TCP sponges were prepared by salt leaching and lyophilization. PLGA 82:18 (Corbion, USA) solution (10% w.v^−1^) was prepared in 1,4-dioxane (Sigma, Germany). β-TCP (Sigma, Germany) was added in it at a concentration of 2.5% w.v^−1^. Sodium chloride (NaCl) particles (diameter range 150-250 µm) were added into the PLGA/TCP suspension in two different concentrations (PLGA/TCP:NaCl = 1:4 or 1:8 w.w^−1^). The mixtures were transferred to Teflon molds (**Figure S1A**), frozen at −80 °C and freeze dried. After leaching the salt in distilled water, sponges were dried at room temperature (RT). The sponges (external dia.10 mm and height 6 mm) had a central cavity (dia. 5 mm and depth 4 mm) for the placement of the collagen-based tumor mimic in (**Figure S1B**).

#### 2.1.2. Preparation of Collagen Sponges (scaffold of the tumor mimic)

Collagen type I was isolated from Sprague-Dawley rat tails as described by earlier studies.(20,21) For the isolation steps, briefly, tendons from rat tails were dissolved in cold acetic acid (0.5 M), filtered, dialyzed against phosphate buffer, and centrifuged. The collagen pellet was dissolved in acetic acid (0.15 M), precipitated with NaCl solution (5% w.v^−1^), dialysis and centrifugation steps were repeated. Collagen precipitate was sterilized in ethanol (70%), frozen at −80 °C and lyophilized. In order to prepare the sponges, collagen solution (1.5 % w.v^−1^ in 0.5 M acetic acid) was placed in the Teflon mold (**Figure S1C**), frozen at −20 °C and lyophilized. They (**Figure S1D**) were then dehydrothermally crosslinked by heating at 140 °C in a vacuum oven.(20,22) Both sponges were characterized before cell seeding. Then the optimized ones were seeded with the required cells and brought together to form BTM.

#### 2.1.3. 3D Bone Tumor Model

The PLGA/TCP and collagen sponges were seeded with the required cells and cultured, and then the cell seeded collagen sponge was inserted in its cavity to form the complete BTM (**Figure S1E**).

### 2.2. Characterization of PLGA/TCP and Collagen Sponges

#### 2.2.1. Contact Angle Measurement

The water contact angle of the sponges was measured using a contact angle goniometer (One Attention, Biolin Scientific, Finland). Distilled water (7 µL) was placed at 5 different locations on the sponges and contact angles were measured by processing the images using the software of the system.

#### 2.2.2. Degradation

Degradation tests of PLGA/TCP sponges were conducted by incubating in sterile phosphate buffer saline (PBS) (0.01 M, pH 7.4) containing sodium azide (0.5 mg.mL^−1^) at 37 °C. Enzymatic degradation of uncrosslinked and DHT-crosslinked collagen sponges were studied in PBS containing collagenase Type II (0.1 mg.mL^−1^, pH 7.4). PBS was replaced at every time point. At each time point, sponges were removed, dried and weighed. Loss of sample weight was calculated by using the equation;

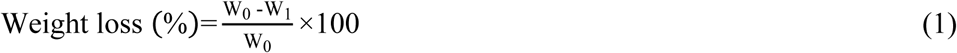

where w_0_ and w_1_ are the dry weights of the samples at time zero and at various time points.

#### 2.2.3. Compressive Mechanical Test

Compression test was performed on the PLGA/TCP and collagen sponges using universal test equipment (Shimadzu AGS-X, Japan, 5 kN load cell) with a compression rate of 0.5 mm.min^−1^.

#### 2.2.4. Scanning Electron Microscopy (SEM)

The samples were placed on carbon tapes (Electron Microscopy Sciences, USA) attached to SEM stubs, and coated with Au–Pd. Micrographs were obtained at 10-20 kV with SEM (QUANTA 400F Field Emission SEM, Netherland) at the METU Central Laboratory.

#### 2.2.4. MicroCT Analysis

The samples were scanned with microcomputed tomography (microCT) (Bruker microCT, SkyScan 1172, Belgium). PLGA/TCP sponges were scanned with application of 100 kV and 100 mA power with Al 0.5 mm filter and collagen sponges by using 35 kV and 21 mA power. Reconstruction was applied with standard software NRecon. Porosity was determined with CTAn software (CTAn, Bruker microCT). Calcium phosphate deposition on the hFOB/HUVEC seeded PLGA/TCP and cell free (control) scaffolds were determined by X-ray absorption spectra using the µCT. Control scaffolds without cells were also cultured in the growth medium for 21 days 34°C in the CO_2_ incubator.

### 2.3. Cell Culture and Seeding

HUVECs (C2517A Lonza, USA) were cultured in EGM-2 BulletKit (Lonza, USA) containing basal medium and SingleQuots™ Kit at 37 °C in a 5% CO_2_ incubator. Human fetal osteoblast cell line, hFOB 1.19 (ATCC, UK), was cultured in DMEM/F12 colorless medium (Gibco, USA) supplemented with fetal bovine serum (FBS) (10%) and G418 (0.3 mg.mL^−1^) (Sigma, Germany) at 34 °C in CO_2_ incubator. Saos-2 cells (ATCC, UK) were cultured in McCoy’s 5A medium (Lonza, USA) supplemented with FBS (15%) and penicillin/streptomycin (100 U mL^−1^) (Sigma, USA) at 37 °C in an incubator. Oxygen plasma was applied to PLGA/TCP (100 W, 3 min) and collagen sponges (100 W, 5 min) to remove the skin layer exist on the surface. Samples were sterilized with UV (30 min for each side). HUVECs were detached with Trypsin-EDTA Solution C and hFOB were detached with Trypsin-EDTA Solution B and cells were collected by centrifugation. Equal number of each cell type (2×10^5^ cells scaffold^−1^) were seeded on the PLGA/TCP scaffolds, and incubated in a 1:1 ratio of DMEM/F12:EGM-2 at 37 °C in an incubator. Saos-2 cells were seeded on collagen scaffolds (1×10^5^ cells scaffold^−1^) and incubated at 37 °C in a CO_2_ incubator. Growth media was refreshed every other day (**Figure 1A, B**).

**Figure 1.**
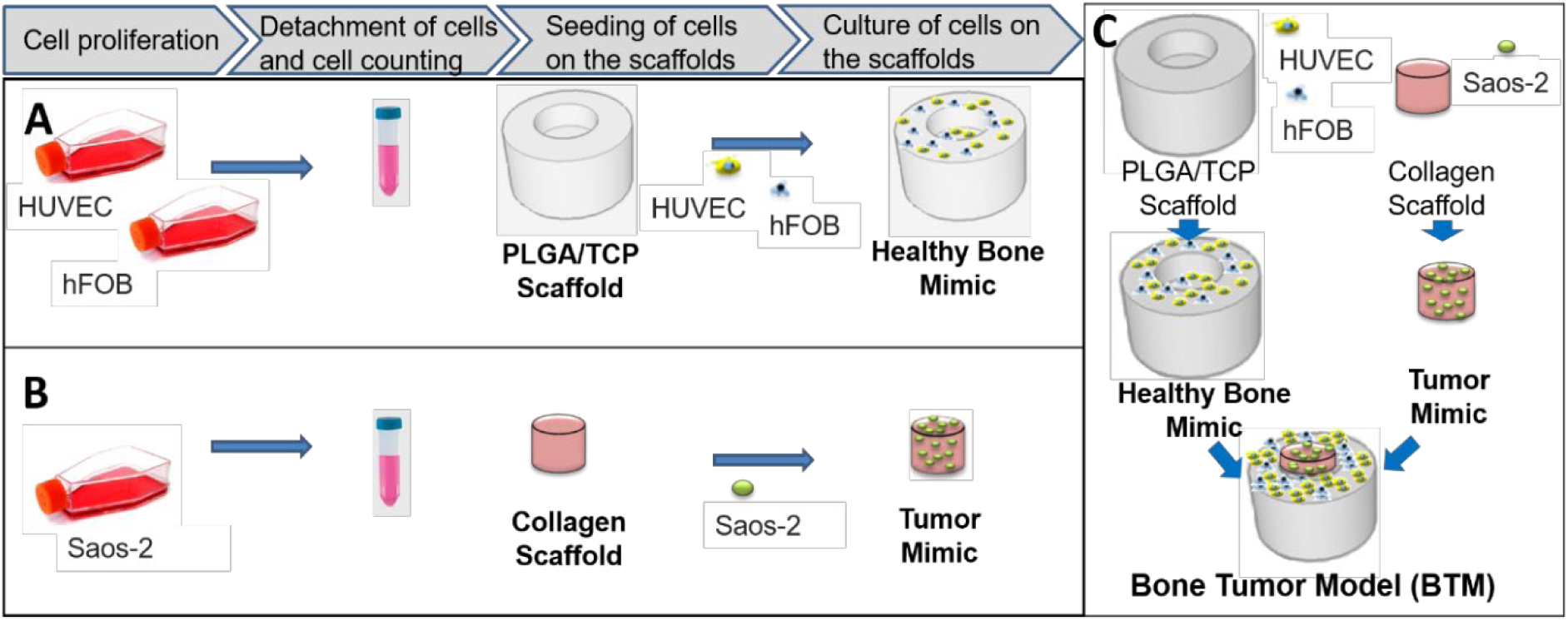
Production of the bone tumor model (BTM). Preparation stages of the (A) healthy bone mimic, and (B) tumor mimic. (C) Integration of tumor mimic in the cavity of the healthy bone mimic to generate bone tumor model (BTM).

### 2.4. Characterization of Healthy Bone and Tumor Mimics

#### 2.4.1. Alamar Blue Cell Viability Assay

Cell seeded scaffolds were washed with PBS and incubated with 10% Alamar Blue solution (Invitrogen, USA) in colorless DMEM for 1 h at 37 °C. The absorbances of the 200 µL solutions were measured at 570 nm and 595 nm with a multiwell plate reader (Molecular Devices, USA). The absorbance value was converted to cell number by using percent reduction values and a calibration curve.

#### 2.4.3. Alkaline Phosphatase (ALP) Activity

SensoLyte pNPP Alkaline Phosphatase Assay Kit was used to determine ALP production by hFOB cells on the PLGA/TCP scaffold. Briefly, samples were washed with component B, lysis buffer was added, frozen and thawed three times at −80 °C and 37 °C. Sonication (50 W, 20 s) was applied and contents were centrifuged (2000 rpm, 10 min). The supernatant (50 µL), ALP dilution buffer (50 µL) and Component A (50 µL) were added in to each well in 96 well plates, incubated for 1 h at 37 °C and then 50 µL stop solution was transferred to each well. Absorbances were measured at 405 nm by a plate reader. ALP concentration was calculated by using the calibration curve.

#### 2.4.3. Immunocytochemistry

The cells present on the scaffolds were fixed by treating them with paraformaldehyde solution (4%) for 15 min at RT. Then, treated with Triton-X-100 solution (1%) for 5 min and incubated in 1% BSA solution at 37 °C to block nonspecific binding. Samples were stained by incubating in Alexa Fluor 532-Phalloidin for 1 h at 37 °C and DRAQ5 for 15 min at RT. Cell seeded samples were also sectioned with cryomicrotome into 20-30 μm thick slices, and transferred to Polysine™ Microscope Adhesion Slides. Sections were also incubated in Triton X-100, blocked in the 1% BSA solution, stained with Alexa Fluor 488 tagged anti-human CD31, Alexa Fluor 532-Phalloidin, and DRAQ5. These samples were examined with a confocal laser scanning microscope (CLSM) (Zeiss LSM 800, Germany).

#### 2.4.4. Scanning Electron Microscopy (SEM)

Cell seeded scaffolds were washed twice with PIPES (piperazine-N,N’-bis(ethanesulfonic acid)) buffer, fixed by immersing the samples in paraformaldehyde (4% w.v^−1^) for 5 min and then lyophilized. Samples were coated with gold-palladium (Au-Pd) under vacuum and examined with SEM (FEI Quanta 200F, USA).

### 2.5. Construction of the Bone Tumor Model (BTM)

hFOB/HUVEC cells were co-cultured on PLGA/TCP scaffolds (as the healthy bone mimic) and Saos-2 cells were cultured on collagen scaffolds (as the tumor mimic) for 2 days. Then, collagen scaffolds were placed in the central cavity of the PLGA/TCP scaffold, and cultured in EGM-2:DMEM/F12: McCoy’s 5A (1:1:1) medium for 21 days (**Figure 1C**).

### 2.6. Characterization of BTM

#### 2.6.1. Immunocytochemistry

Immunostaining was conducted as described in Section 2.4.3 on the bicomponent construct. Cryosections were incubated in 1X blocking solution (5% goat serum, 1% Tween 20, 1% BSA and 1% sodium azide in PBS) for 1h at 37 °C, incubated in anti-CD31 primary Ab (10 µg mL^−1^ in 0.1X blocking solution) and anti-von Willebrand Factor (vWF) primary Ab (0.05 µg.mL^−1^ in 0.1X blocking solution) at 4 °C overnight, washed with PBS and incubated in donkey anti-sheep secondary Ab (Alexa Fluor® 594) (0.02 µg.mL^−1^ in 0.1X blocking solution) and goat anti-rabbit secondary Ab (Alexa Fluor® 647) (0.02 µg.mL^−1^ in 0.1X blocking solution) for 1 h at 37 °C. Samples were washed with PBS, incubated in Alexa Fluor 488-Phalloidin for 1 h at 37 °C, and in DAPI for 15 min at RT and examined with CLSM.

#### 2.6.2. Quantitative Real-Time PCR (qRT-PCR) Analysis of Angiogenesis

Angiogenesis was studied using qRT-PCR (Rotor-Gene Q; Qiagen) and 2^−ΔΔCt^ relative quantification method. Primers were synthesized for vascular endothelial growth factor (VEGF), basic fibroblast growth factor (bFGF), and interleukin 8 (IL-8) genes by Sentegen (Sentegen, Turkey) according to the sequences are presented in **Table 1**. Three types of samples were analyzed: Saos-2 on the control, TCPS, Saos-2 on collagen scaffolds alone (coded as Coll), and Saos-2 in collagen scaffolds placed in the cavity of PLGA/TCP scaffolds co-cultured hFOB/HUVEC (coded as Coll/BTM). On days 7, 14 and 21, RNA was isolated using the RNeasy Micro Kit (Qiagen) according to the manufacturer’s protocol. The isolated RNA samples were treated with DNase I (DNA-free™ Kit; Ambion, Invitrogen, Germany) to prevent DNA contamination and converted to cDNA using RevertAid First Strand cDNA Synthesis Kit (Invitrogen). qRT-PCR was performed with the cDNA samples obtained.

**Table 1.**
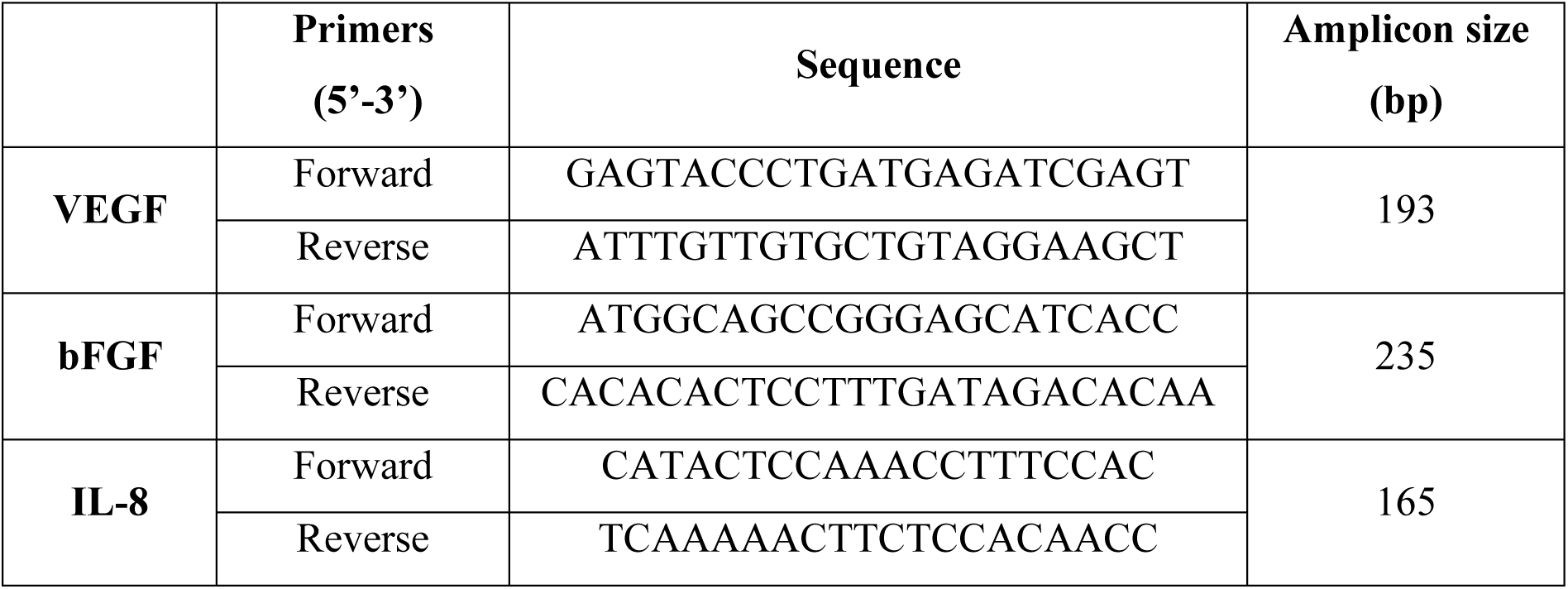
Quantitative Real-Time PCR Primers and Amplicon Sizes.

#### 2.6.3. Agarose Gel Electrophoresis

The agarose gel (2% in Tris-EDTA buffer solution) was placed in the separator buffer tank, into which the electrodes are placed. Samples were loaded into the wells and 100 V potential was applied across the electrodes. Samples were run on the gel for 75 min and visualized under UV light (UVP GelDoc Imaging System, USA).

### 2.7. Efficacy of Anticancer Agent Tested on the BTM

BTM was developed as described previously and cultured in EGM-2: DMEM/F12: McCoy’s 5A (1:1:1) medium for 7 days. Then, the samples were incubated in medium containing Doxorubicin (2.7 µg.mL^−1^ Doxorubicin) for 24 h. The medium was refreshed and incubated for another 24 h. After that, the following efficacy studies were carried out for both the drug-treated samples and the drug-free controls.

#### 2.7.1. Cell Number by Alamar Blue Assay

Alamar Blue cell viability assay was performed on the samples by modifying previous Alamar Blue assay procedure. After incubation with the drug for 24 h, the samples were incubated for 3 days in the fresh drug-free medium, 10% Alamar Blue solution in colorless DMEM was added on the samples and then incubated for a further 3 h at 37 °C and 5% CO_2_ condition. The absorbance of 200 µL solution was measured at 570 nm and 595 nm with a plate reader. Live/dead analyses of the cells were determined by CLSM. For this purpose, the samples were washed with PBS and stained with calcein AM (0.5 µL.mL^−1^ in PBS) for live cells, and ethidium homodimer-1 (2 µL.mL^−1^ in PBS) for dead cells.

#### 2.7.2. Determination of Caspase-3 Activity

Caspase-3 activity was determined using Caspase-3 Activity Kit (Fluorometric) (Abcam, USA) on Doxorubicin treated BTM according to the manufacturer’s protocol. Cell lysis buffer was added on the sample and incubated on ice for 10 min. DTT and DEVD-AFC enzyme substrates were added and the whole solution was incubated for 2 h at 37 °C. Fluorescence intensities at 400 nm (λ_ex_) and 505 nm (λ_em_) were measured with a plate reader.

### 2.8. Statistical Analysis

All quantitative data in this study were expressed as mean ± standard deviations with n≥3 unless otherwise stated. Statistical analysis was performed with GraphPad Prism6 program. Differences between group means were analyzed with Student’s T-test when the data were normally distributed. Comparisons of groups with one independent variable were performed with One-way ANOVA with Tukey’s posthoc test, to determine significant differences. When there were two independent variables, Two-way ANOVA was used. All values are represented as the mean ± standard deviation. Differences of p < 0.05 were considered significant.

## 3. Results and Discussion

We prepared a 3D-BTM which mimics the bone tumor together with health microenvironment around the tumor. Each compartment of the BTM examined separately to optimize the preparation conditions and then combined to form the final tissue model. Healthy and tumor parts were prepared from PLGA/TCP and collagen. PLGA/β-TCP scaffolds were used in many studies and demonstrated good cell-material interaction and osteoconductivity,(23) enhancement in the bone regeneration of critical bone defects,(24) better guidance in the culture of osteoblasts(25) and improvement in biological activity, such as calcium deposition.(26) Thus, β-TCP containing PLGA scaffolds were chosen to present the healthy component of the model. As the tumor matrix, collagen sponges were selected because collagen constitutes 95% of the organic part of the bone matrix and it is a widely used biomaterial for 3D modeling due to its biocompatibility, biodegradability, crosslinking capacity.(27)

### 3.1. Properties of PLGA/TCP Scaffolds

Healthy bone mimics were prepared by freeze drying and salt leaching methods using two different NaCl content with PLGA and TCP together and coded as PLGA:TCP:NaCl (4:1:20) and PLGA:TCP:NaCl (4:1:40), where the numbers define weight ratios. The average pore size of both scaffolds was about 199±52 µm as determined by semi-quantitative image analysis (n ≥ 20) mode of ImageJ (NIH, USA), and porosities determined from the microCT images were 96.7% and 92.6% for the samples PLGA:TCP:NaCl (4:1:40) and PLGA:TCP:NaCl (4:1:20), respectively. Results prove that existing of higher porogen content (NaCl in our case) had higher porosity in accordance with the literature.(28,29) These results were also supported by SEM and microCT images (**Figure 2A, B**). SEM also showed that treatment of the samples with O_2_ plasma removed the ‘skin layer’ that generally forms on the surfaces during lyophilization and blocks the openings of the pores (**Figure 2C**).(30,31). O_2_ plasma treated PLGA/TCP sponges were used in cell seeding and culture experiments because they enhanced hydrophilicity and had open pores on the surface as a result of the removal of the skin layer by plasma treatment. O_2_ plasma also helped enhance hydrophilicity of the surface a shown by the decrease in the contact angle of the PLGA/TCP scaffold from 114.5°±8.7° to 100.0°±6.1° (**Figure 2D**). It is reported that anchorage-dependent mammalian cells favor moderately hydrophilic surfaces.(32-34) We also have shown that cells do not prefer highly hydrophilic nor hydrophobic surfaces to attach.(35,36) Thus the surfaces we obtained were not optimal but they were more suitable for cell seeding than the pristine PLGA/TCP form. Degradation studies showed that the scaffolds are very stable and lost only about 2% of their weight in 7 weeks incubation in PBS (**Figure 2E**). These results also show the suitability of the PLGA scaffolds for long term in vitro drug screening tests.(37) Compressive elastic modulus of PLGA:TCP:NaCl (4:1:20) and PLGA:TCP:NaCl (4:1:40) were determined as 5.18 MPa and 4.76 MPa, respectively, showing that the higher porogen (NaCl) content, increased the porosity and led to a decrease in the mechanical strength (**Figure 2F**). Similar results were also reported by He et al. for six cylindrical PLLA sponges with different NaCl content. They also reported that higher NaCl particle content led to a higher porosity and lower mechanical strength.(28) It is known that high porosity and pore interconnectivity are essential for cell migration and proliferation, as well as for proper nutrient exchange in the culture medium. The best scaffolds would be those with high and interconnected porosity, and high mechanical strength. Since the difference between mechanical strengths of the two scaffolds tested was not very high, the sponge with higher porosity, PLGA:TCP:NaCl (4:1:40), was selected.

**Figure 2.**
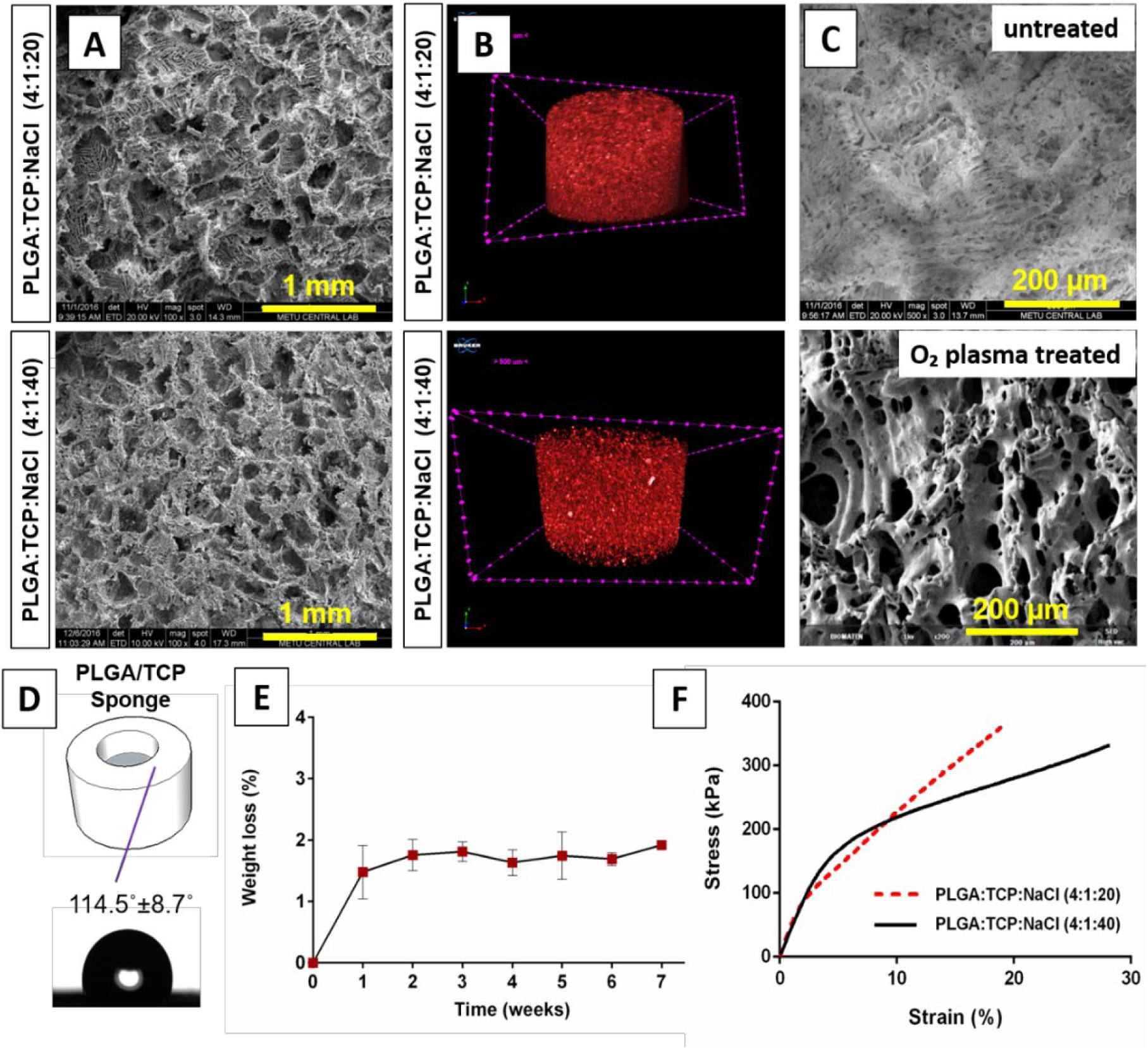
Characterization of cell-free PLGA/TCP sponges. (A) SEM micrographs of the horizontal cross section, and (B) MicroCT images of the sponges PLGA:TCP:NaCl (4:1:20) (upper row), PLGA:TCP:NaCl (4:1:40) (lower row). (C) SEM micrographs of pristine and oxygen plasma-treated PLGA/TCP sponge surfaces. (D) Contact angle (n=5), (E) degradation behavior (n=3) (one-way ANOVA followed by Tukey post-hoc test, results were significant), and (F) compression test results of both PLGA/TCP sponges (n=6).

### 3.2. Characterization of Collagen Sponges

Porous collagen sponges prepared by lyophilization were examined with SEM and microCT. Horizontal and longitudinal cross sections of these sponges were very similar without any significant difference (**Figure 3A**). All sections had high porosity and interconnectivity which is required for efficient transport of nutrients and waste.(38) Porosities of the sponges were determined from both SEM and µCT data as 96.7% and 86% for PLGA:TCP:NaCl (4:1:40) and collagen sponges, respectively (**Figure 3B, C**). The average diameters of the pores were measured by using ImageJ (NIH, USA) as 199±52 µm for PLGA:TCP:NaCl (4:1:40) and 50-150 μm for collagen scaffolds.

**Figure 3.**
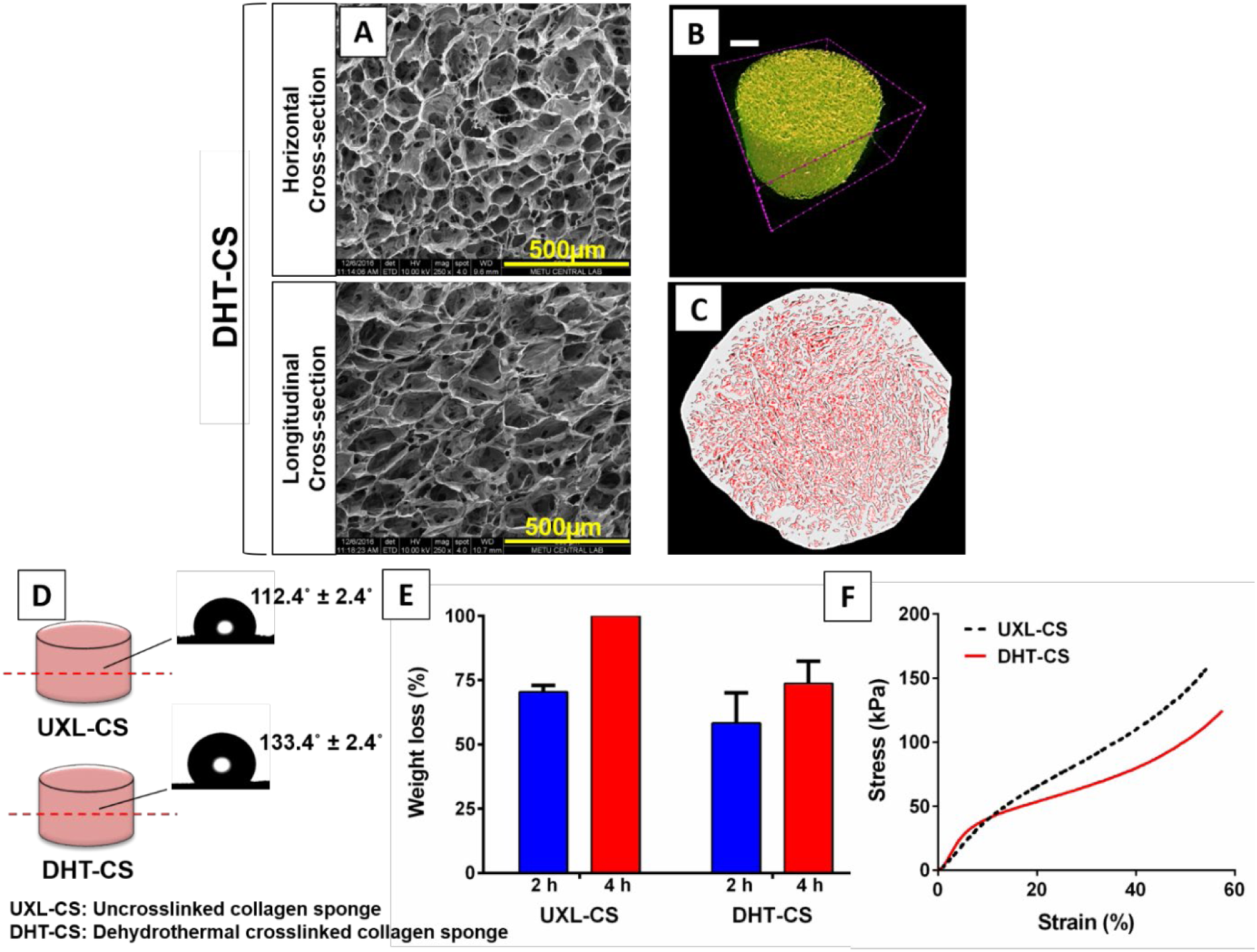
Characterization of dehydrothermal crosslinked collagen sponges (DHT-CS) and uncrosslinked collagen sponges (UXL-CS). (A) SEM micrographs of the horizontal and longitudinal cross section of DHT-CS. MicroCT images of the DHT-CS: (B) side view, and (C) top view. Scale bar: 250 μm. (D) Contact angles (n=5) (Student’s T-test was used, result were significant), (E) enzymatic degradation test (n=3) (Student’s T-test was used, result were significant), and (F) compressive mechanical test results of the UXL-CS and DHT-CS (n=6).

In bone tissue engineering, scaffolds are usually produced with pore sizes similar to that of trabecular bone (20-1500 μm).(38) It was also reported that pores in the range 160–270 μm support rapid and extensive angiogenesis within a scaffold.(39) The pore size of our PLGA:TCP constructs (199±52 μm) which were used to grow hFOB and HUVEC cells is in the range of the ideal pore size of the bone matrix. On the other hand, the osteoblasts were shown to populate more in smaller pores (40 μm) when they were grown in scaffolds with different pore sizes; larger pore sizes (100 μm) facilitated cell migration. Also, the minimum porosity necessary for the regeneration of a blood vessel was reported as 30-40 μm to enable the exchange of metabolic components and to facilitate endothelial cell entrance and 10-100 μm to allow cell infiltration.(38) As a result, the pore size of the collagen scaffold (50-150 μm) used in our study is suitable for cell growth, infiltration and migration.

The contact angle of uncrosslinked collagen sponge (UXL-CS) and dehydrothermal crosslinked collagen sponge (DHT-CS) were found to be 112.4° ± 2.4° and 133.3° ± 2.2°, respectively (**Figure 3D**). Results show that the sponges became more hydrophobic after crosslinking possibly because during dehydrothermal treatment, ester and amide bonds are created either by esterification or amide formation decreasing the amount of the free carboxyl, amine, and hydroxyl groups and result in more hydrophobic materials lead to the formation of crosslinking between triple helix chains of collagen via condensation reactions.(40)

In the enzymatic stability tests, collagenase is used and it breaks the peptide bonds of collagen and cause destruction of extracellular matrix.(41) After 2 h incubation in collagenase solution, the weight loss for UXL-CS (70.5% ± 2.5) was higher than DHT-CS (58.3% ± 11.8). When we extended the treatment from 2 h to 4 h, UXL-CS completely disintegrated while DHT-CS lost a large fraction (73.8% ± 8.6) of its initial weight (**Figure 3E**). A higher degree of stability (and therefore degradation) of DHT-CS was expected due to the crosslinking treatment.

Compressive elastic modulus of UXL-CS and DHT-CS were 111±18 kPa and 140±46 kPa as measured from the initial slopes of stress-strain curves. This shows that crosslinking enhanced the mechanical strength of the sponges in addition to increasing its stability (**Figure 3F**). Based on these results, DHT-CS was selected as the matrix to culture Saos2 cells and to serve as the tumor mimic.

### 3.3. Properties of Tumor and Healthy Bone Mimics

#### 3.3.1. Cell Proliferation and ALP Activity

Alamar Blue assay was performed on hFOB/HUVEC cells (1:1) seeded on the PLGA/TCP scaffold to study the cell proliferation (**Figure 4A**). Cell seeding density was 2×10^5^ cells and on Day 1 approximately 2.7×10^5^ cells were counted on the TCPS control while much less (8.7×10^4^) cells were determined on the PLGA/TCP scaffold. Cell number on the scaffold increased gradually over time while on TCPS, the cell number rapidly reached a plateau in a week indicating confluence.

**Figure 4.**
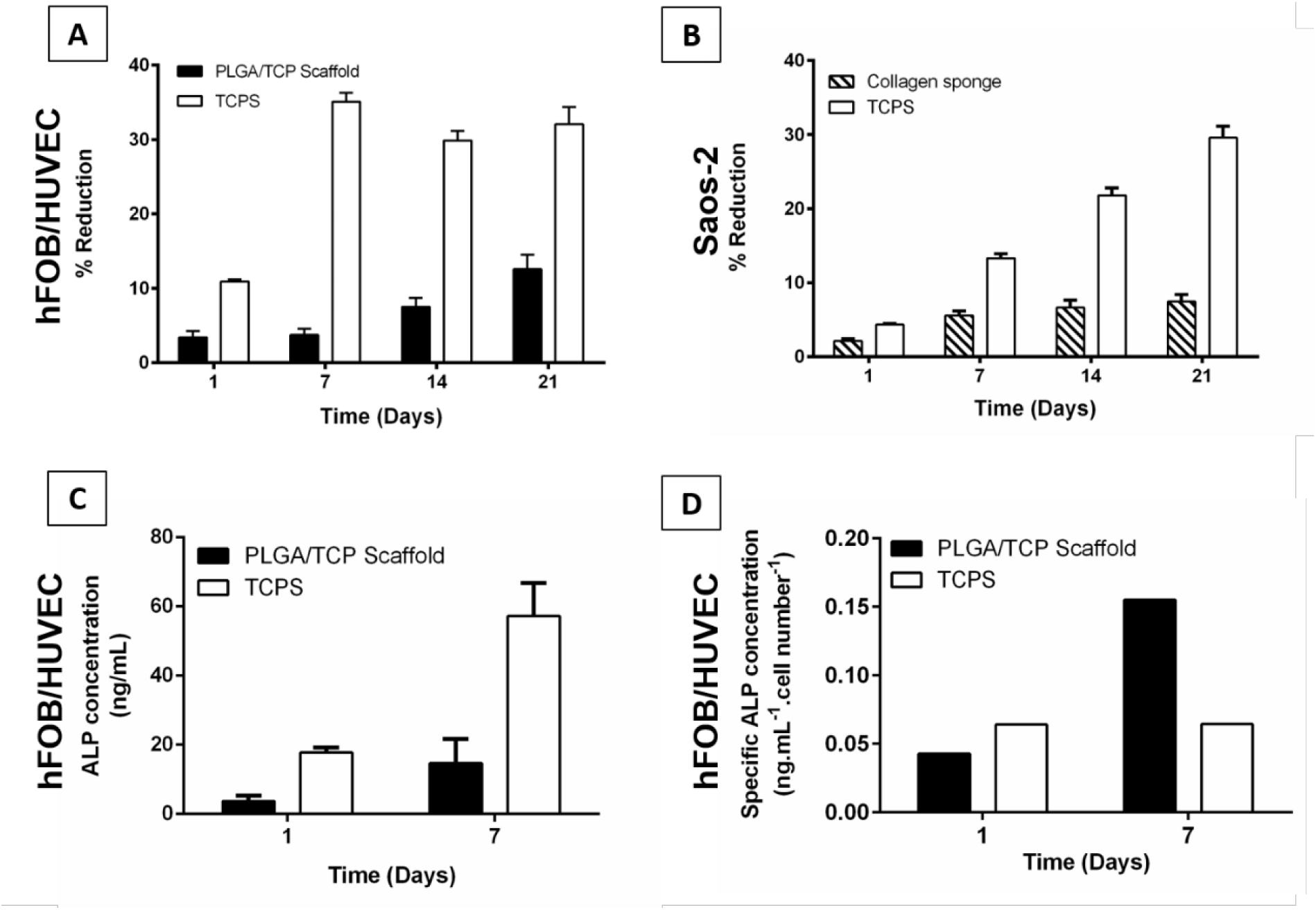
Cell proliferation and ALP activity on cell seeded PLGA and collagen scaffolds. (A) Proliferation of hFOB/HUVEC on PLGA/TCP scaffolds. (B) Proliferation of Saos-2 on the collagen sponge. (C) Alkaline phosphatase (ALP) activity of hFOB cells on PLGA/TCP scaffolds. (D) Specific ALP activity of the hFOB on PLGA/TCP scaffolds normalized to cell number obtained from Alamar Blue cell viability assay. Seeding densities: hFOB/HUVEC: 2×10^5^ scaffold^−1^ and Saos-2: 1×10^5^ scaffold^−1^. (A, B, C: two-way ANOVA followed by Sidak’ multiple comparison test, n=3, p<0.0001, results were significant).

Alamar Blue assay was also performed for Saos-2 cells seeded on the collagen scaffold (**Figure 4B)**. Cell seeding density was half of that of the hFOB/HUVEC (1×10^5^). Rate of proliferation of Saos-2 on TCPS was higher than on the collagen as was on the PLGA/TCP scaffold. In this case, however, the cell number increase on TCPS continued for 3 weeks but on collagen, the rate of cell number increase and the cell number itself were lower than on TCPS.

ALP activity of hFOB on PLGA/TCP scaffold was determined on Days 1 and 7 (**Figure 4C**). It increased during the 7 day incubation; however, ALP activity on the scaffold was about 3 times lower than the control. Since the cell numbers were not the same on different scaffolds the ALP values were normalized by dividing the results with the cell numbers. The normalized ALP activity on Day 7 was significantly (about 2-fold) higher (**Figure 4D**) showing that the cells on the sponges could produce significant amounts of ALP.

#### 3.3.2. Microscopy and MicroCT Results

Nuclei and cytoskeletons of the cells were stained to study cell morphology, intercellular interaction, and cell-material interactions using CLSM and SEM. From Day 7 onwards, the cells spread over the surface and covered it completely (**Figure 5A**). They showed filamentous extensions (filopodia) indicating interaction with the surface (**Figure 5B**). Anti-human CD31 immunostaining of HUVECs on PLGA/TCP scaffolds (**Figure 5C**) showed expression of CD31. Since this protein is excreted in high levels by early and mature endothelial cells, it is used to show the presence of HUVECs on the specimen (42). The CD31 staining in our samples proves the presence of the HUVECs which were seeded on the scaffolds. Staining of actin and nuclei also shows that both HUVECs and hFOBs exist on the section, which were also seeded on the scaffolds. Calcium phosphate (CaP) crystals produced by cells in addition to the TCP particles introduced during scaffold production are observed as white spots on the microCT images (**Figure 5D**). However, it is not possible to distinguish those produced by the cells and those were introduced during scaffold production from each other.

**Figure 5.**
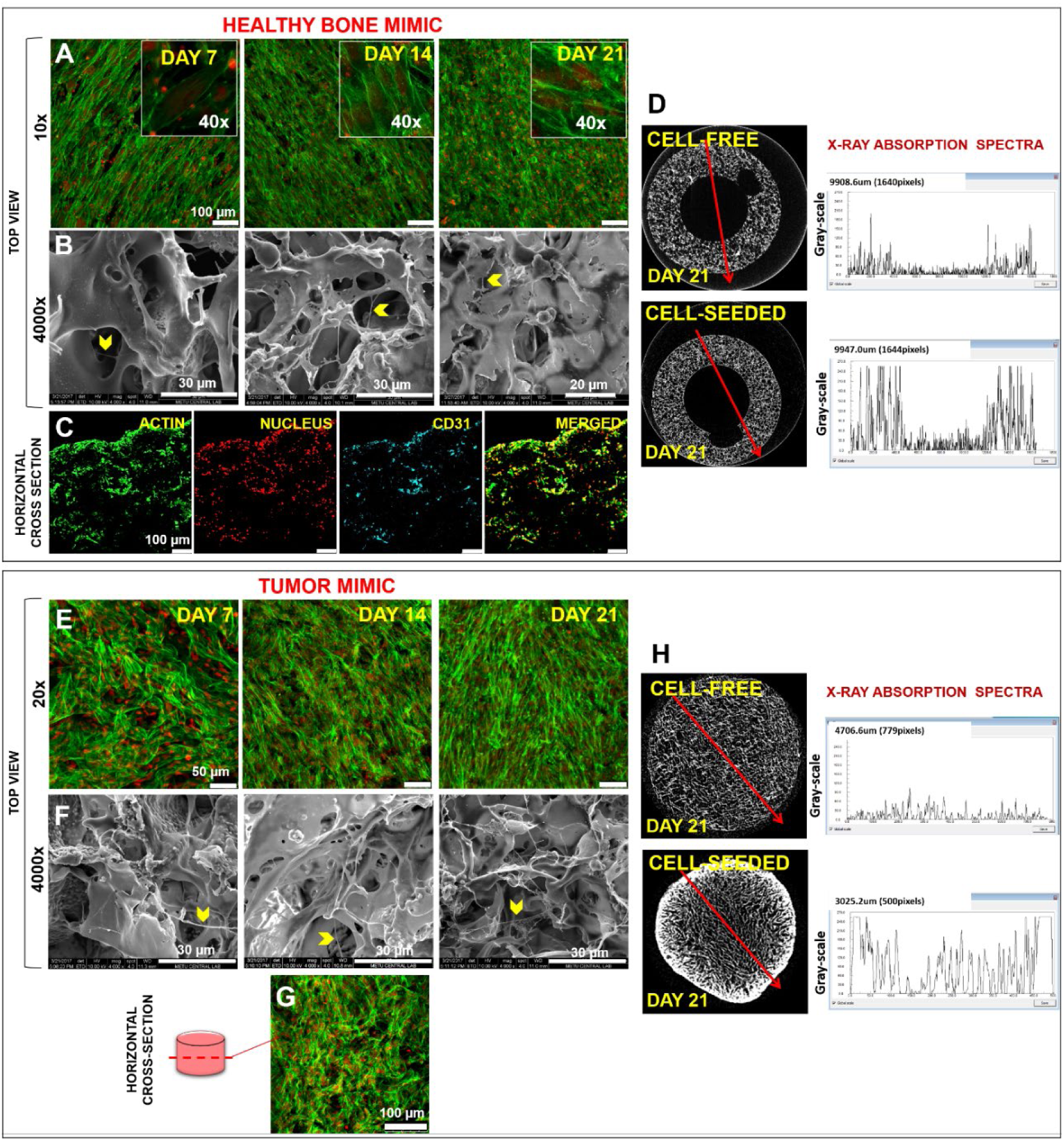
CLSM, SEM and microCT analysis results. (A) CLSM and (B) SEM micrographs of hFOB and HUVECs on PLGA/TCP scaffold on Days 7, 14 and 21. (C) CLSM micrographs of the horizontal cross section of PLGA/TCP scaffolds on Day 28. (D) MicroCT images (Top View) and X-ray absorption spectra of the horizontal cross sectional of the healthy bone mimic. (E) CLSM, and (F) SEM micrographs of the Saos-2 on collagen scaffolds on Days 7, 14 and 21. (G) CLSM micrograph of a horizontal cross section of the collagen scaffold. (H) MicroCT images (Top View) and X-ray absorption spectra of the horizontal cross sectional views of tumor mimic. Cell seeding densities= 1×10^5^ Saos-2 scaffold^−1^ and 2×10^5^ hFOB+HUVEC scaffold^−1^. The stains were Alexa Fluor 532 Phalloidin (green): actin, DRAQ5 (red): nucleus and Alexa Fluor 488 anti-human CD31 (blue):CD31. Yellow arrowheads indicate filamentous cell extensions (filopodia). The red lines show the direction in which the spectra were taken.

Changes in porosity were observed upon cell seeding and the culture duration. The 96.7% porosity of unseeded PLGA/TCP scaffolds decreased with time to 89.6, 90.2 and 85.7% on Days 7, 14 and 21, respectively. This decrease in the porosity is probably because of the filling of the pores by the production of the new extracellular matrix and deposition of new calcium phosphate by the cells.(43) The peaks on the X-ray absorption spectra of hFOB/HUVEC seeded PLGA/TCP scaffolds were also higher than the cell-free scaffolds **(Figure 5D, right)** and are most probably due to the calcium phosphate deposition by cells. The CLSM results of Saos-2 cells on the collagen tumor mimic are shown in **Figure 5E-H**. The cells started to spread, fill the pores of the collagen sponge, and interact with each other as the incubation time increased (**Figure 5E**). On Days 14 and 21, the coverage is more extensive and the cells appear to be organized in the form of a layer. These increases were also indicated by the Alamar blue assay of the Saos-2 cells (**Figure 4B**). Here, the cell number did not increase further possibly because of the confluence of the cells. Cells interact with the surface with their filopodia (**Figure 5F**). Meanwhile, it is also seen that the cells had migrated and attached to the walls of the pores at the core of the sponge (**Figure 5G**). MicroCT analysis shows calcium phosphate forming capacities of the cells (**Figure 5H**).

Cell presence, proliferation and secretion of calcium phosphate affected the porosity of the collagen sponges as they did with the PLGA scaffolds. Porosity of cell-free collagen sponge was 86%, and decreased to 73% and 56% after 7 days and 3 weeks of culturing, respectively. When microCT images of horizontal cross section of the unseeded and cell seeded sponges were investigated, the exterior of the collagen sponge where cells proliferated and formed cell clusters appeared brighter than the core of the sponges (**Figure 5H**). X-ray absorption of the unseeded sample (examined along the red line) is almost linear demonstrating the homogenous composition of the collagen sponge (**Figure 5H, right**).

### 3.4. Properties of the Complete Bone Tumor Model (BTM)

#### 3.4.1. Microscopy Results

The complete bone tumor model (BTM) has two compartments and is formed by combination of tumor mimic and healthy bone mimic. Experiments carried out on individual sponges (PLGA and collagen) were also performed with the BTM. Cell nuclei and cytoskeleton were stained and since the dyes stained the cells indiscriminately, the cells could not be distinguished from each other **(Figure 6A-D)**. Cells on the BTM attached, spread and covered the surfaces of the scaffolds by Day 21. **In Figure 6E-H**, samples were stained with anti-von Willebrand factor (vWF) (red) and CD31 antibody (pink) dyes, both of which are specific for HUVECs. Cytoskeleton (green) and nucleus (blue) of the cells were also stained. Due to the specificity of these dyes, it became possible to distinguish HUVECs from the other two cell types, and they were detected on the exterior and the top surface of the collagen sponge (**Figure 6E, F**) in addition to exterior and bottom surfaces of the PLGA/TCP scaffold (**Figure 6G, H**). Since HUVECs and hFOB were seeded only on the PLGA/TCP scaffolds, the presence of HUVECs on the PLGA/TCP scaffold was expected, but their detection on the collagen sponge demonstrates that these cells have migrated from the healthy bone mimic to the tumor mimic area. This migration of the HUVECs to the cancer component was an indication of the angiogenic process which precludes the vascularization of the tumor tissue as in the native biological systems.(44) Spreading and coverage of the surfaces of sponges by hFOB, HUVECs and Saos-2 was also shown by the SEM micrographs (**Figure 6I-L**). Horizontal sections of the BTM (**Figure 6M, N**) were stained to show the migration of HUVECs to the core of the cancer mimic (collagen). Cells of the healthy bone mimic were mainly detected in the regions that are in direct contact with the collagen core, the cancer mimic. HUVECs (pink) that have migrated into the tumor area are clearly seen in **Figure 6N** (magnified image of Figure 6M). The direct contact between the scaffolds apparently allowed the migration of the endothelial cells to the tumor side as if to vascularize the tumor. This is a good indicator that the model works. The microCT of the horizontal section of BTM also showed this direct contact between the two regions indicating that this model has a good potential to mimic the cancer tissue (**Figure 6O**).

**Figure 6.**
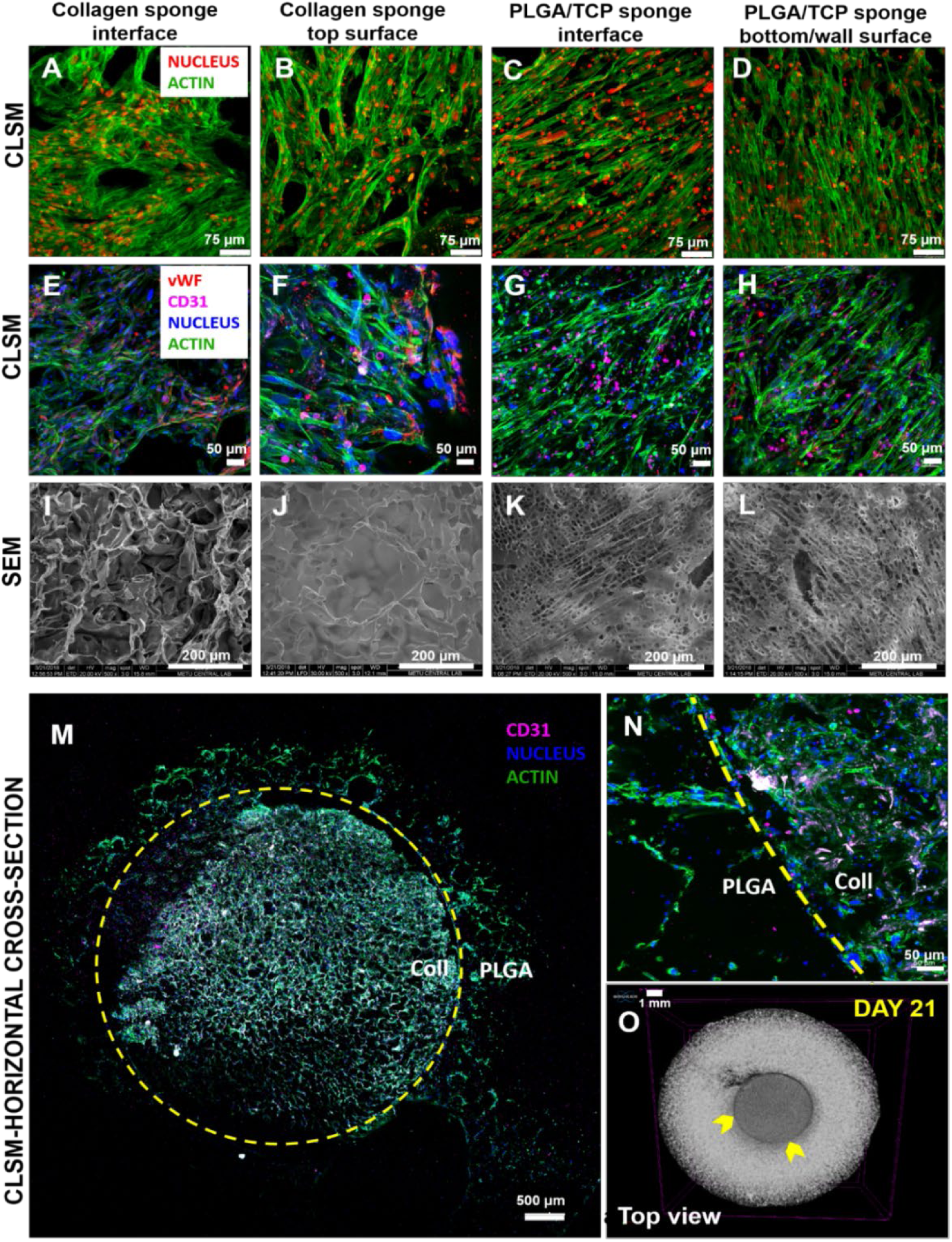
CLSM and SEM analysis of BTM. (A-D) CLSM micrographs of actin and nucleus staining, (E-H) antibody staining and (I-L) SEM micrographs of hFOB, HUVECs, and Saos-2 on PLGA/TCP and collagen scaffolds on Day 21. Alexa Fluor 532 Phalloidin (green) stains actin and DRAQ5 (red) stains nucleus of all type of cells in Figure A-D. Alexa Fluor 488 Phalloidin (green) stains actin, DAPI (blue) stains nucleus of all type of cells and anti-CD31 antibody (pink) and anti-vWF antibody (red) stains HUVECs in Figure E-H. (M, N) CLSM micrographs, and (O) microCT image of a horizontal section of BTM. Yellow arrow-heads and yellow, dashed lines show the interface between the two mimics. Alexa Fluor 488 Phalloidin (green) stains actin, DAPI (blue) stains nucleus of all type of cells. Anti-CD31 antibody (pink) stains HUVECs.

#### 3.4.2. Molecular Analysis of Angiogenesis

Expression of angiogenic factors (VEGF, bFGF, and IL-8) in the BTM was studied with real-time quantitative PCR (RT-PCR) for 21 days (**Figure 7**). In this analysis, three types of samples were used: Saos-2 cells cultured on: 1) TCPS (control group), 2) collagen sponge, and 3) collagen sponge in BTM.

**Figure 7.**
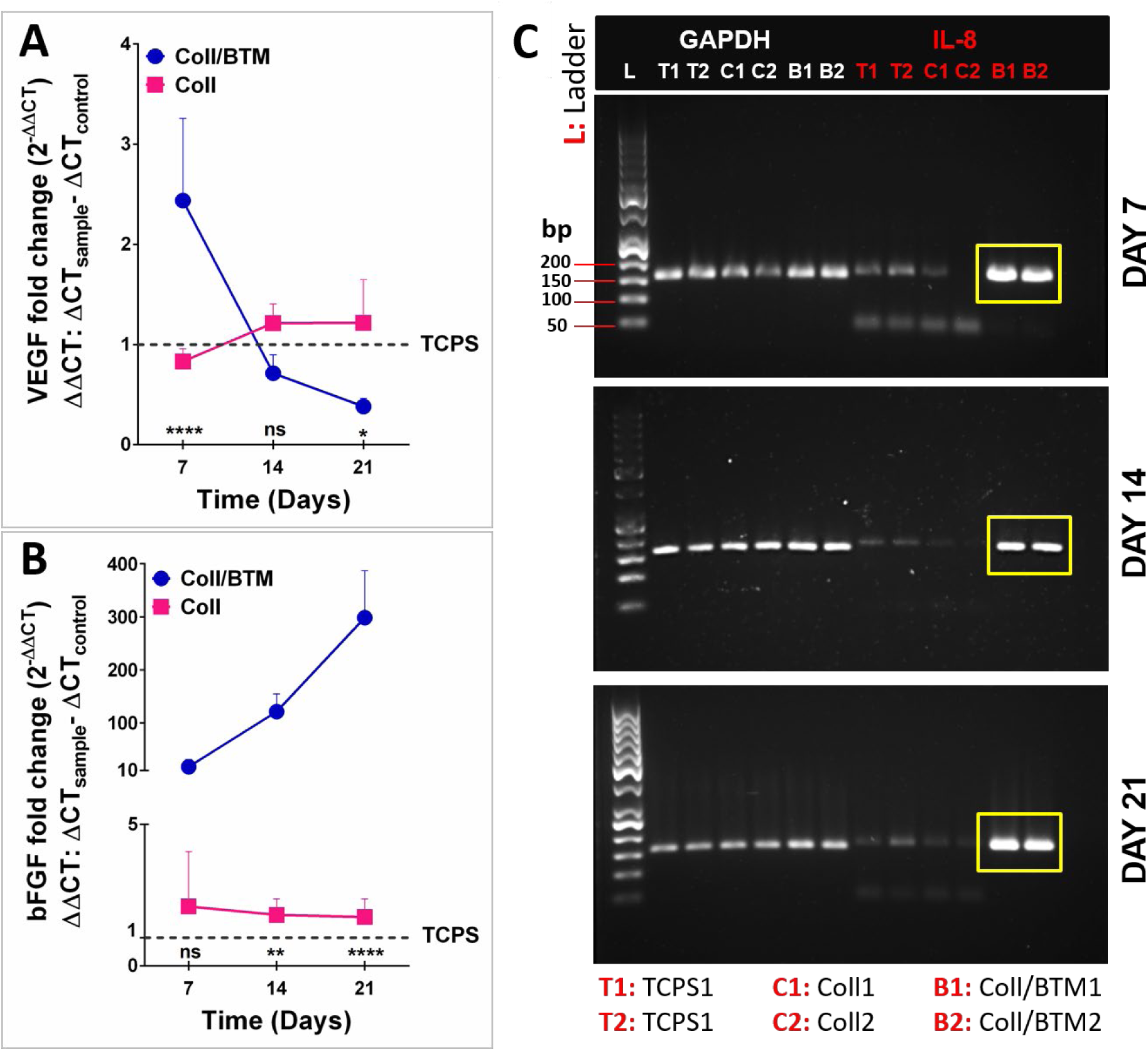
Analysis of angiogenesis of the BTM. Relative expression levels of (A) VEGF and (B) bFGF genes, and (C) agarose gel electrophoresis of IL-8 genes of Saos-2 cells on TCPS surface (TCPS), collagen sponge (Coll) and collagen sponge in bone tumor model (Coll/BTM) on Days 7, 14 and 21. Yellow rectangles show the cDNA bands corresponding to IL-8 mRNA of Saos-2 in Coll/BTM. Statistical analysis between samples (Coll and Coll/BTM) was carried out using two-way ANOVA. *p < 0.05, **p < 0.01, ****p < 0.0001, and ns: not significant, n=4.

At certain time points, the collagen sponges were treated for RNA isolation and qRT-PCR analysis. In case of BMT model, the collagen sponges were separated from PLGA/TCP scaffolds. Relative VEGF and bFGF gene expression levels are presented in **Figure 7A, B**. Gene expression level of Saos-2 cells cultured on TCPS served as reference. No significant differences were detected in the VEGF expression on Days 7, 14 and 21 by Saos-2 cells cultured in collagen sponges. On the other hand, in BTM, VEGF expression by Saos-2 was significantly high on Day 7. The literature states that the presence of the healthy bone mimic around the tumor tissue affects the expression level of VEGF, and that of tumor progression as was observed in our study.(45,46) VEGF is a powerful angiogenic factor, and it is generally believed that tumor cells secrete VEGF to enhance the formation of its own vascular network.(47) It stimulates the formation of new blood vessels and controls apoptosis and differentiation of tumor cells and osteoblasts. It has an effect on tumor progression and pathological remodeling. However, the VEGF expression level in the later days decreased probably as a result of a negative feedback mechanism. As reported in some studies,(48-50) there are some negative regulators that influence angiogenesis in an autocrine manner leading to downregulation of VEGF. Some of these regulators are directly induced by stimulators of angiogenesis, especially VEGF, as a consequence of a specific negative-feedback regulator mechanism of angiogenesis.(51) There was also no significant difference in bFGF expression by Saos-2 cells in the collagen sponge alone, but in case of BTM these values were higher by approximately 10, 110 and 300 times on Days 7, 14 and 21, respectively. These high levels and the substantial increase are probably a synergic effect of the interactions and cross-talk between the hFOB and HUVECs seeded into the model.

IL-8 has some critical effects on pro-angiogenic, pro-migratory, and osteoclastogenic activities. Many cancerous cell types express IL-8 resulting in proliferation and migration of cancer cells and tumor angiogenesis and metastasis, and researchers have shown that highly metastatic solid tumors express IL-8.(52) In this study, relative IL-8 gene expressions of the samples could not be calculated because the expression levels by Saos-2 on TCPS surface and collagen sponge was too low for quantification with qRT-PCR. For this reason, the IL-8 gene expression in BTM was demonstrated by agarose gel electrophoresis. Micrographs show significantly high levels of IL-8 expression in the BTM on Days 7, 14 and 21 (**Figure 7C**). Tan et al. co-cultured U2OS osteosarcoma cells with immortalized fibroblasts or HUVECs to study the effect of 3D structure (silk scaffolds) and tumor–stroma interaction and reported that the co-culture of U2OS osteosarcoma cells with immortalized fibroblasts resulted in the upregulation of angiogenic factors (VEGF, IL-8, bFGF) and especially that of IL-8.(53) In the present study, co-culture of Saos-2, in close vicinity of hFOB and HUVECs, significantly upregulated the IL-8 expression. As a result, we showed that the expression of proangiogenic factors increased significantly in the two compartment novel bone tumor model, indicating the success of mimicking the natural tissue.

### 3.5. Study of Efficacy of the Anticancer Agent on Cell Viability on BTM

In order to assess the suitability of the constructed BTM to serve as a cancer tissue model, Doxorubicin (a drug used in osteosarcoma therapy) was added into the system and its effectiveness on viability of the cells was studied. Initially, a dose-response curve was prepared in vitro with Saos-2 cells in the tissue culture plate and exposed to a range of concentrations of the drug. Dose-Response curve (**Figure S2**) showed that the IC_50_ value of Doxorubicin was 0.1876 µg.mL^−1^ when the seeding density was 2×10^4^ cells per well. Since almost the cells on the BTM was 15 fold higher than on the TCPS wells (3×10^5^ cells), 15 fold higher drug concentration (2.7 µg.mL^−1^ Doxorubicin) was used on the model. Alamar Blue cell viability test showed that the cell number was significantly decreased following treatment with Doxorubicin (**Figure 8A**). Caspase-3 enzyme activity is an indicator of apoptosis and **Figure 8B** shows the activity (thus the amount) of the caspase-3 enzyme in the Doxorubicin-administered BTM. Doxorubicin was applied to the bicomponent bone tumor model, and then the two cell-seeded sponges (exterior PLGA and core collagen) were separated from each other for the analysis of the enzymatic activity on each component. A significant increase in caspase-3 activity was determined in the collagen sponge, while there was no significant change in healthy bone tissue mimic. It was expected that the drug would be more effective on cancer tissue mimic because of the cancer cells loaded in it. Live/Dead assay also showed lower viability in the Doxorubicin-treated model than the untreated one (**Figure 8C**). Doxorubicin was more effective at the top surface of the collagen sponge possibly because this part of the model was in more direct contact with the drug containing culture medium because it is exposed to it (**Figure 8C, top row**). A drug dose killing all the cells in the tissue culture plates but only a portion in the BTM is also a good indicator of the difference in the effect of the drugs on 2D and 3D cultures. Apparently, the cells in the culture plate (the 2D environment) are more exposed to the culture conditions while in the 3D BTM model the microenvironment prevents such direct effects. In **Figure 8C**, the collagen sponge bottom surface and the PLGA/TCP sponge cavity bottom surface are not in direct contact with the drug-containing medium and so the number of affected cells is lower than those at the top surface of the collagen sponge. It can, therefore, be concluded that the BTM developed in this study mimics the bone tumor; of course, further experiments are needed to optimize the parameters before initiating any patient specific human trials.

**Figure 8.**
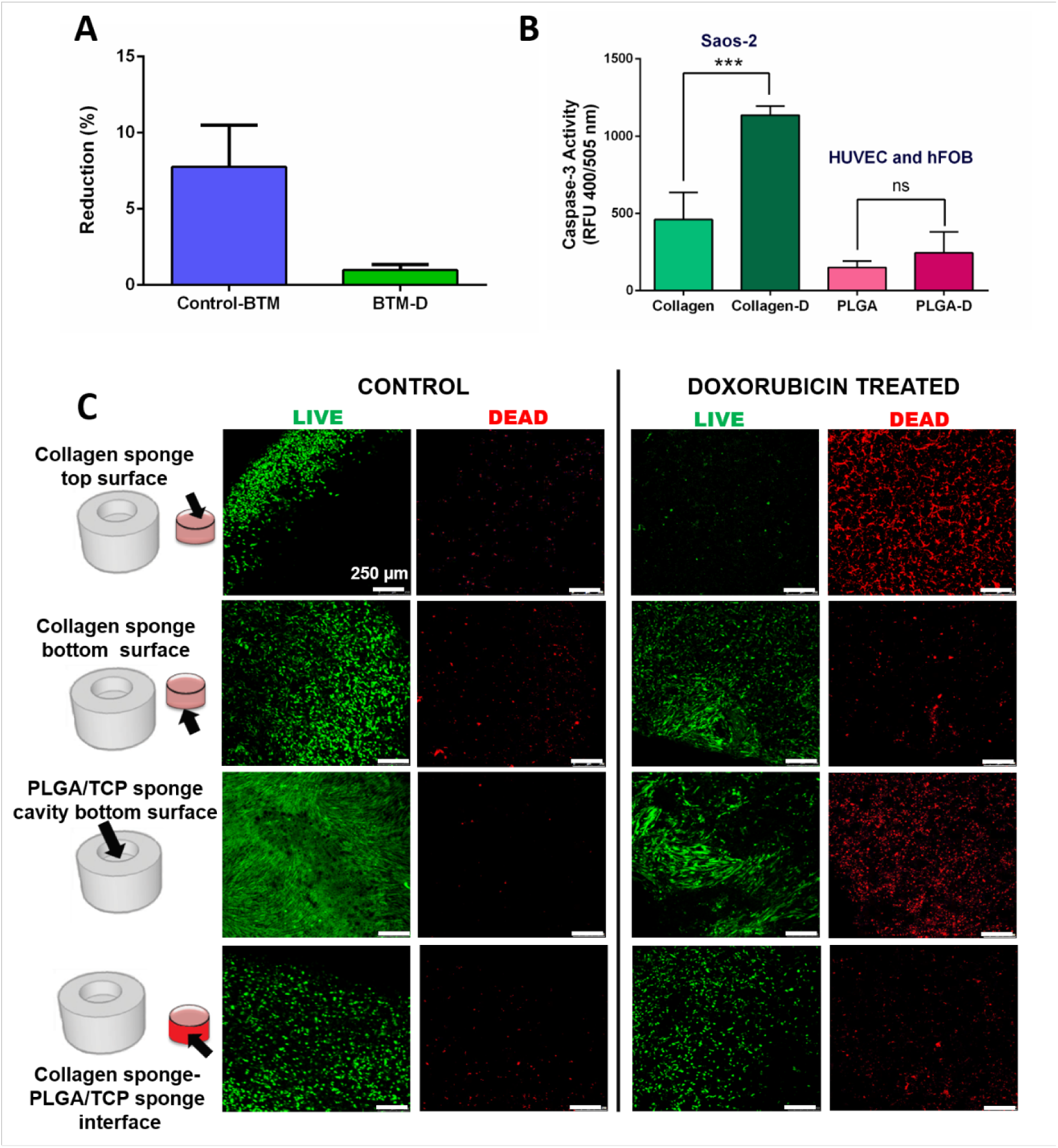
Efficacy of Doxorubicin on the BTM. (A) Alamar Blue cell viability assay of Doxorubicin-treated (BTM-D) and untreated BTM (Control-BTM). Reduction (%) of the dye is proportional to live cell number. (B) Caspase-3 enzyme activity. (C) Cell viability assay of Doxorubicin-treated and untreated BTM using Live/Dead staining. CLSM micrographs of stained cells on different parts of the model are given. Calcein AM (green) shows viable cells, ethidium homodimer-1 (red) shows nonviable cells. Scale bar= 250 µm for all images. (A: Student’s T-test was used, n=3, results were significant, B: one-way ANOVA followed by Tukey post-hoc test, result were significant ***p < 0.001, and ns: not significant, n=3).

The results presented in the study showed that the bone tumor model could mimic the natural osteosarcoma and would be considered for studying the efficacy of different types of drugs on cells obtained from a patient for a ‘patient specific treatment’. However, several optimization steps should be performed. Although proliferation of Saos-2 cells and their interactions with HUVECs were observed, formation of spheroids was not. In the literature, there are studies carried out with spheroids. It is known that tumor cells in spheroids have strong cell-cell interactions and a different morphology than cells in a 2D monolayer culture. Interactions between cells in 3D spheroids also increase their stability and survival rates showing resistance to anti-cancer agents.(54-56) Thus, studies with spheroids need to be performed and the core collagen should carry these instead of individual cells. Studies along this line are in progress.”

## 4. Conclusion

In this study, a functional 3D bone tumor model (BTM) was successfully developed by combining two components: 1) a tumor mimic consisting of osteosarcoma (Saos-2) cells seeded in a collagen sponge, and 2) healthy bone mimic that was formed by seeding hFOB and HUVECs in PLGA/TCP sponge. This model aimed to mimick the *in vivo* tumor stroma by providing not only the cross-talk between the cancer cells, but also the interactions of cancer cells with the healthy bone cells and the bone matrix. The natural bone tumor microenvironment constructed in this manner led to cancer cells increasing their expression of angiogenic factors when cultured in the BTM model. A direct contact between the healthy bone cells (hFOB and HUVECs) and cancer cells (Saos-2) was achieved, and when treated with the anticancer agent, Doxorubicin, the effect was observed specifically on the cancer cells. These indicate that this new strategy in tumor modeling, with tumor cells cultured within an engineered bone construct, could create a microenvironment similar to that of the native tissue. In conclusion, we believe the 3D BTM developed in this study could be a good candidate for screening tests of drug efficacy. This will also be a promising approach in the personalized drug therapy in finding the right type, combination and dose regimens of the drugs for a specific individual.

## Supporting Information

Photographs of Teflon molds and the sponges and dose response curve of Saos-2 cells grown in a tissue culture plate treated with Doxorubicin.

## Acknowledgements

Authors thank TUBITAK for the support through Project No: 215M621 and the Grant BIDEB2211 – C, and A.K. acknowledges the scholarships through the same. Authors also thank BIOMATEN, METU Center of Excellence in Biomaterials and Tissue Engineering for the support and the use of the facilities.

## References

1. Misaghi, A.; Goldin, A.; Awad, M.; Kulidjian, A. A. Osteosarcoma: a comprehensive review. SICOT-J 2018, 4, 12, DOI: 10.1051/sicotj/2017028.

2. Abarrategi, A.; Tornin, J.; Martinez-Cruzado, L.; Hamilton, A.; Martinez-Campos, E.; Rodrigo, J. P.; González, M.V.; Baldini, N.; Garcia-Castro, J.; Rodriguez, R. Osteosarcoma: cells-of-origin, cancer stem cells, and targeted therapies. Stem Cells Int. 2016, 2016, 1–13, DOI: 10.1155/2016/3631764.

3. Verrecchia, F.; Rédini, F. Transforming growth factor-β signaling plays a pivotal role in the interplay between osteosarcoma cells and their microenvironment. Front. Oncol. 2018, 8, 133, DOI: 10.3389/fonc.2018.00133.

4. Botter, S. M.; Neri, D.; Fuchs, B. Recent advances in osteosarcoma. Curr. Opin. Pharmacol. 2014, 16, 15–23, DOI: 10.1016/J.COPH.2014.02.002.

5. Alemany-Ribes, M.; Semino, C.E. Bioengineering 3D environments for cancer models. Adv. Drug Deliv. Rev. 2014, 79, 40–49, DOI: 10.1016/j.addr.2014.06.004.

6. Duval, K.; Grover, H.; Han, L.-H.; Mou, Y.; Pegoraro, A. F.; Fredberg, J.; Chen, Z. Modeling Physiological Events in 2D vs. 3D Cell Culture. Physiology 2017, 32, 266–277, DOI: 10.1152/physiol.00036.2016.

7. Pampaloni, F.; Reynaud, E. G.; Stelzer, E. H. K. The third dimension bridges the gap between cell culture and live tissue. Nat. Rev. Mol. Cell Biol. 2007, 8, 839–845, DOI: 10.1038/nrm2236.

8. Kapalczynska, M.; Kolenda, T.; Przybyła, W.; Zajączkowska, M.; Teresiak, A.; Filas, V. 2D and 3D cell cultures – a comparison of different types of cancer cell cultures, Arch. Med. Sci. 2018, 14, 910–919, DOI: 10.5114/aoms.2016.63743.

9. Lama, R.; Zhang, L.; Naim, J. M.; Williams, J.; Zhou, A.; Su, B. Development, validation and pilot screening of an in vitro multi-cellular three-dimensional cancer spheroid assay for anti-cancer drug testing. Med. Chem. 2013, 21, 922–931, DOI: 10.1016/j.bmc.2012.12.007.

10. Lamhamedi-Cherradi, S. E.; Santoro, M.; Ramammoorthy, V.; Menegaz, B. A.; Bartholomeusz, G.; Iles, L. R.; Amin, H. M.; Livingston, J. A.; Mikos, A. G.; Ludwig, J. A. 3D tissue-engineered model of Ewing’s sarcoma. Adv. Drug Deliv. Rev. 2014, 79–80, 155–171, DOI: 10.1016/j.addr.2014.07.012.

11. Charoen, K. M.; Fallica, B.; Colson, Y. L.; Zaman, M. H.; Grinstaff, M. W. Embedded multicellular spheroids as a biomimetic 3D cancer model for evaluating drug and drug-device combinations. Biomaterials 2014, 35, 2264–2271, DOI: 10.1016/j.biomaterials.2013.11.038.

12. Martins-Neves, S. R.; Lopes, Á. O.; do Carmo, A.; Paiva, A. A.; Simões, P. C.; Abrunhosa, A. J.; Gomes, C.M. Therapeutic implications of an enriched cancer stem-like cell population in a human osteosarcoma cell line. BMC Cancer. 2012, 12, 139, DOI: 10.1186/1471-2407-12-139.

13. Baek, N.; Seo, O. W.; Kim, M.; Hulme, J.; An S. S. A. Monitoring the effects of doxorubicin on 3D-spheroid tumor cells in real-time. Onco. Targets. Ther. 2016, 9, 7207–7218, DOI: 10.2147/OTT.S112566.

14. Fong, E. L. S.; Lamhamedi-Cherradi, S.-E.; Burdett, E.; Ramamoorthy, V.; Lazar, A. J.; Kasper, F. K.; Farach-Carson, M. C.; Vishwamitra, D.; Demicco, E. G.; Menegaz, B. A.; Amin, H. M.; Mikos, A. G.; Ludwig, J. A. Modeling Ewing sarcoma tumors in vitro with 3D scaffolds. Proc. Natl. Acad. Sci. U.S.A. 2013, 110, 6500–6505, DOI: 10.1073/pnas.1221403110.

15. Tan, P. H. S.; Aung, K. Z.; Toh, S. L.; Goh, J. C. H.; Nathan, S. S. Three-dimensional porous silk tumor constructs in the approximation of in vivo osteosarcoma physiology. Biomaterials 2011, 32, 6131–6137, DOI: 10.1002/term.1800.

16. Fallica, B.; Maffei, J. S.; Villa, S.; Makin, G.; Zaman, M. Alteration of Cellular Behavior and Response to PI3K Pathway Inhibition by Culture in 3D Collagen Gels. PLoS One. 2012, 7, e48024, DOI: 10.1371/journal.pone.0048024.

17. Villasante, A.; Marturano-Kruik, A.; Vunjak-Novakovic, G. Bioengineered human tumor within a bone niche. Biomaterials. 2014, 35, 5785–5794, DOI: 10.1016/j.biomaterials.2014.03.081.

18. Villasante, A.; Marturano-Kruik, A.; Robinson, S. T.; Liu, Z.; Guo, X. E.; Vunjak-Novakovic, G. Tissue-engineered model of human osteolytic bone tumor. Tissue Engineering Part C: Methods. 2017, 23, 98–107, DOI: 10.1089/ten.TEC.2016.0371.

19. Schuessler, T. K.; Chan, X. Y.; Chen, H. J.; Ji, K.; Park, K. M.; Roshan-Ghias, A.; Kuhn, N. Z. Biomimetic tissue-engineered systems for advancing cancer research: NCI Strategic Workshop report. Cancer Research. 2014, 74, 5359–5363, doi: 10.1158/0008-5472.CAN-14-1706.

20. Kilic, C.; Girotti, A.; Rodriguez-Cabello, J. C.; Hasirci, V. A collagen-based corneal stroma substitute with micro-designed architecture. Biomater. Sci. 2014, 2, 318–329, DOI: 10.1039/C3BM60194C.

21. Komez, A.; Baran, E. T.; Erdem, U.; Hasirci, N.; Hasirci, V. Construction of a patterned hydrogel-fibrous mat bilayer structure to mimic choroid and Bruch’s membrane layers of retina. J. Biomed. Mater. Res. Part A. 2016, 104, 2166–2177, DOI: 10.1002/jbm.a.35756.

22. Drexler, J. W.; Powell, H. M. Dehydrothermal crosslinking of electrospun collagen. Tissue Eng. Part C Methods 2011, 17, 9–17, DOI: 10.1089/ten.tec.2009.0754.

23. Pang L, Hu Y, Yan Y.; Liu, L.; Xiong, Z.; Wei, Y.; Bai, J. Surface modification of PLGA/β-TCP scaffold for bone tissue engineering: Hybridization with collagen and apatite. Surf Coatings Technol. 2007, 201(24), 9549–9557, DOI:10.1016/j.surfcoat.2007.04.035

24. Lin, L.; Gao, H. Modification of β-TCP/PLGA scaffold and its effect on bone regeneration in vivo. J Wuhan Univ Technol Sci Ed. 2016, 31(2), 454–460, DOI:10.1007/s11595-016-1391-y

25. Yang, Y.; Tang, G.; Zhao, Y.; Zhang, Y.; Li, X.; Yuan, X. Effect of degradation of PLGA and PLGA/β-TCP scaffolds on the growth of osteoblasts. Chinese Sci Bull. 2011, 56(10), 982–986, DOI:10.1007/s11434-010-4132-1

26. Kim, S. H.; Kim, S. H.; Jung, Y. Bi-layered PLCL/(PLGA/β-TCP) composite scaffold for osteochondral tissue engineering. J Bioact Compat Polym. 2015, 30(2), 178–187, DOI:10.1177/0883911514566015

27. Fitzgerald, K. A.; Guo, J.; Tierney, E. G.; Curtin, C. M.; Malhotra, M.; Darcy, R.; O’Brien, F. J.; O’Driscoll, C. M. The use of collagen-based scaffolds to simulate prostate cancer bone metastases with potential for evaluating delivery of nanoparticulate gene therapeutics. Biomaterials 2015, 66, 53–66, DOI:10.1016/j.biomaterials.2015.07.019

28. He, X.; Kawazoe, N.; Chen, G. Preparation of cylinder-shaped porous sponges of poly(L-lactic acid), poly(DL-lactic-co-glycolic acid), and poly(ε-caprolactone). Biomed Res. Int. 2014, 2014, 1–8, DOI: 10.1155/2014/106082.

29. Iannace, S.; Maio, E.; Di Nicolais, L. Preparation and Characterization of Polyurethane Porous Membranes by Particulate-leaching Method. Cellular Polymers 2001, 20, 321–338, DOI: 10.1177/026248930102000502.

30. Lehocký, M.; Drnovská, H.; Lapcíková, B.; Barros-Timmons, A.; Trindade, T.; Zembala, M.; Lapcík, L. Plasma surface modification of polyethylene. Colloids and Surfaces A: Physicochem. Eng. Aspects 2003, 222, 125–131, DOI: 10.1016/S0927-7757(03)00242-5.

31. Cvelbar, U.; Mozetic, M.; Klanjsek-Gunde, M. Selective oxygen plasma etching of coatings. IEEE T. Plasma Sci. 2005, 33, 236–237. DOI: 10.1109/TPS.2005.845345.

32. Nagahara, S.; Matsuda, T. Cell-substrate and cell-cell interactions differently regulate cytoskeletal and extracellular matrix protein gene expression. J. Biomed. Mater. Res. 1996, 32, 677–686, DOI: 10.1002/(SICI)1097-4636(199612)32:4&lt;677::AID-JBM22&gt;3.0.CO;2-9.

33. Gumbiner, B. M. Cell adhesion: the molecular basis of tissue architecture and morphogenesis. Cell 1996, 84, 345–357.

34. Ozcan, C.; Hasirci, N. Plasma modification of PMMA films: surface free energy and cell-attachment studies. J. Biomater. Sci. Polym. Ed. 2007, 18, 759–773, DOI: 10.1163/156856207781034124.

35. Hasirci, V.; Tezcaner, A.; Hasirci, N.; Süzer, S. Oxygen plasma modification of poly(3-hydroxybutyrate-co-3-hydroxyvalerate) film surfaces for tissue engineering purposes. J. Appl. Polym. Sci. 2003, 87, 1285–1289, DOI: 10.1002/app.11532.

36. Vardar, E.; Endogan, T.; Kiziltay, A.; Hasirci, V.; Hasirci, N. Effect of oxygen plasma on surface properties and biocompatibility of PLGA films. Surface and Interface Analysis 2010, 42, 486–491, DOI: 10.1109/BIYOMUT.2010.5479756.

37. Lu, L.; Peter, S. J.; Lyman, M. D.; Lai, H.-L.; Leite, S. M.; Tamada, J. A.; Uyama, S.; Vacanti, J. P.; Langer, R.; Mikos, A. G. In vitro and in vivo degradation of porous poly(dl-lactic-co-glycolic acid) foams. Biomaterials 2000, 21, 1837–1845, DOI: 10.1016/S0142-9612(00)00047-8.

38. Loh, Q. L.; Choong, C. Three-dimensional scaffolds for tissue engineering applications: role of porosity and pore size. Tissue Eng. Part B Rev. 2013, 19, 485–502, DOI: 10.1089/ten.TEB.2012.0437.

39. Artel, A.; Mehdizadeh, H.; Chiu. Y-C; Brey E. M.; Cinar, A. An Agent-Based Model for the Investigation of Neovascularization Within Porous Scaffolds. Tissue Eng Part A. 2011, 17(17-18), 2133–2141, DOI:10.1089/ten.tea.2010.0571

40. Pulieri, E.; Chiono, V.; Ciardelli, G.; Vozzi, G.; Ahluwalia, A.; Domenici, C.; Vozzi, F.; Giusti, P. Chitosan/gelatin blends for biomedical applications. J. Biomed. Mater. Res. Part A. 2008, 86, 311–322, DOI: 10.1002/jbm.a.31492.

41. Chung, L.; Dinakarpandian, D.; Yoshida, N.; Lauer-Fields, J. L.; Fields, G. B.; Visse, R. Nagase, H. Collagenase unwinds triple-helical collagen prior to peptide bond hydrolysis. The EMBO Journal. 2004, 23, 3020–3030, DOI: 10.1038/sj.emboj.7600318.

42. Edwards, N.; Langford-Smith, A. W. W.; Wilkinson, F. L.; Alexander, M. Y. Endothelial progenitor cells: new targets for therapeutics for inflammatory conditions with high cardiovascular risk. Front. Med. 2018, 5, 200, DOI: 10.3389/fmed.2018.00200.

43. Intranuovo, F.; Howard, D.; White, L. J.; Johal, R. K.; Ghaemmaghami, A. M.; Favia, P.; Howdle, S. M.; Shakesheff, K. M.; Alexander, M. R. Uniform cell colonization of porous 3-D scaffolds achieved using radial control of surface chemistry. Acta Biomater. 2011, 7, 3336–3344, DOI: 10.1016/J.ACTBIO.2011.05.020.

44. Chaddad, H.; Kuchler-Bopp, S.; Fuhrmann, G.; Gegout, H.; Ubeaud-Sequier, G.; Schwinté, P.; Bornert, F.; Benkirane-Jessel, N.; Idoux-Gillet, Y. Combining 2D angiogenesis and 3D osteosarcoma microtissues to improve vascularization. Experimental Cell Research 2017, 360, 138–145, DOI: 10.1016/j.yexcr.2017.08.035.

45. Bachelder, R. E.; Crago, A.; Chung, J.; Wendt, M. A.; Shaw, L. M.; Robinson, G.; Mercurio, A. M. Vascular endothelial growth factor is an autocrine survival factor for neuropilin-expressing breast carcinoma cells. Cancer Res. 2001, 61, 5736–5740.

46. Deckers, M. M. L.; Karperien, M.; van der Bent, C.; Yamashita, T.; Papapoulos, S. E.; Löwik, C. W. Expression of vascular endothelial growth factors and their receptors during osteoblast differentiation. Endocrinology 2000, 141, 1667–1674, DOI: 10.1210/endo.141.5.7458.

47. Peng, N.; Gao, S.; Guo, X.; Wang, G.; Cheng, C.; Li, M.; Liu, K. Silencing of VEGF inhibits human osteosarcoma angiogenesis and promotes cell apoptosis via VEGF/PI3K/AKT signaling pathway. J. Transl. Res. 2016, 8, 1005–1015.

48. Coch, L.; Mejias, M.; Berzigotti, A.; Garcia-Pras, E.; Gallego, J.; Bosch, J.; Mendez, R. Fernandez, M. Disruption of negative feedback loop between vasohibin-1 and vascular endothelial growth factor decreases portal pressure, angiogenesis, and fibrosis in cirrhotic rats. Hepatology 2014, 60, 633–647, DOI: 10.1002/hep.26995.

49. Lobov, I. B.; Renard, R. A.; Papadopoulos, N.; Gale, N. W.; Thurston, G.; Yancopoulos, G. D.; Wiegand, S. J. Delta-like ligand 4 (Dll4) is induced by VEGF as a negative regulator of angiogenic sprouting. Proc. Natl. Acad. Sci. 2007, 104, 3219–3224, DOI: 10.1073/pnas.0611206104.

50. Suzuki, Y.; Kobayashi, M.; Miyashita, H.; Ohta, H.; Sonoda, H.; Sato, Y. Isolation of a small vasohibin-binding protein (SVBP) and its role in vasohibin secretion. J. Cell Sci. 2010, 123, 3094–3101, DOI: 10.1242/jcs.067538.

51. Saito, T.; Takeda, N.; Amiya, E.; Nakao, T.; Abe, H.; Semba, H.; Soma, K.; Koyama, K.; Hosoya, Y.; Imai, Y.; Isagawa, T.; Watanabe, M.; Manabe, I.; Komuro, I.; Nagai, R.; Maemura, K. VEGF-A induces its negative regulator, soluble form of VEGFR-1, by modulating its alternative splicing. FEBS Lett. 2013, 587, 2179–2185, DOI: 10.1016/J.FEBSLET.2013.05.038.

52. Ning, Y.; Manegold, P. C.; Hong, Y. K.; Zhang, W.; Pohl, A.; Lurje, G.; Winder, T.; Yang, D.; La Bonte, M. J.; Wilson, P. M.; Ladner, R. D.; Lenz, H. J.; Interleukin-8 is associated with proliferation, migration, angiogenesis and chemosensitivity in vitro and in vivo in colon cancer cell line models. Int. J. cancer. 2011, 128, 2038–2049, DOI: 10.1002/ijc.25562.

53. Tan, P. H. S.; Chia, S. S.; Toh, S. L.; Goh, J. C. H. Nathan, S. S. The dominant role of IL-8 as an angiogenic driver in a three-dimensional physiological tumor construct for drug testing. Tissue Eng. Part A. 2014, 20, 1758–1766, DOI: 10.1089/ten.TEA.2013.0245.

54. Zanoni, M.; Piccinini, F.; Arienti, C. 3D tumor spheroid models for in vitro therapeutic screening: a systematic approach to enhance the biological relevance of data obtained. Sci Rep. 2016, 6, 19103, DOI:10.1038/srep19103

55. Charoen, K. M.; Fallica, B.; Colson, Y. L.; Zaman, M. H.; Grinstaff, M. W. Embedded multicellular spheroids as a biomimetic 3D cancer model for evaluating drug and drug-device combinations. Biomaterials 2014, 35(7), 2264–2271. DOI:10.1016/j.biomaterials.2013.11.038

56. Lama, R.; Zhang, L.; Naim, J. M.; Williams, J.; Zhou, A.; Su, B. Development, validation and pilot screening of an in vitro multi-cellular three-dimensional cancer spheroid assay for anti-cancer drug testing. Bioorg Med Chem. 2013, 21(4), 922–931. DOI:10.1016/j.bmc.2012.12.007

