## Supporting Information for "A 2-Compartment Bone Tumor Model for Testing the Efficacy of Cancer Drugs"

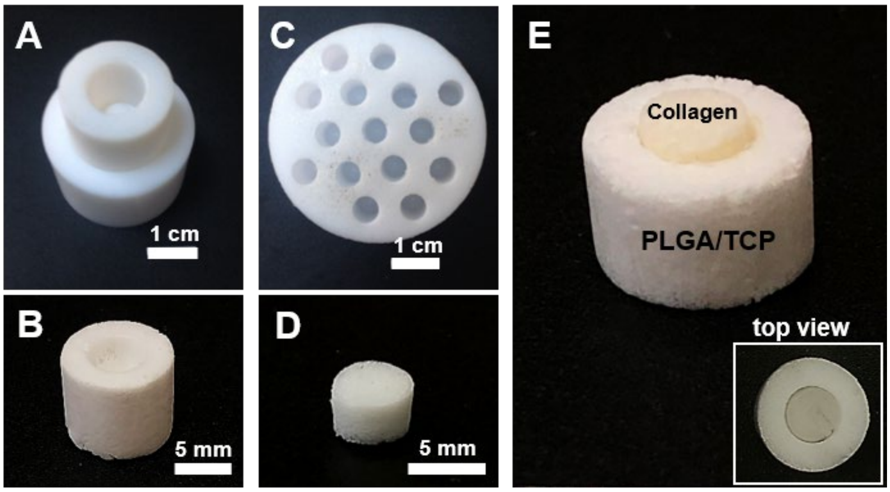


**Figure S1.** Photographs of Teflon molds and the sponges. Photographs of Teflon molds for preparation of (A) PLGA/TCP sponges, and (C) collagen sponges. Photographs of (B) PLGA/TCP sponge, (D) collagen sponge and (E) the complete model with collagen sponges placed in the cavity of PLGA/TCP sponge.

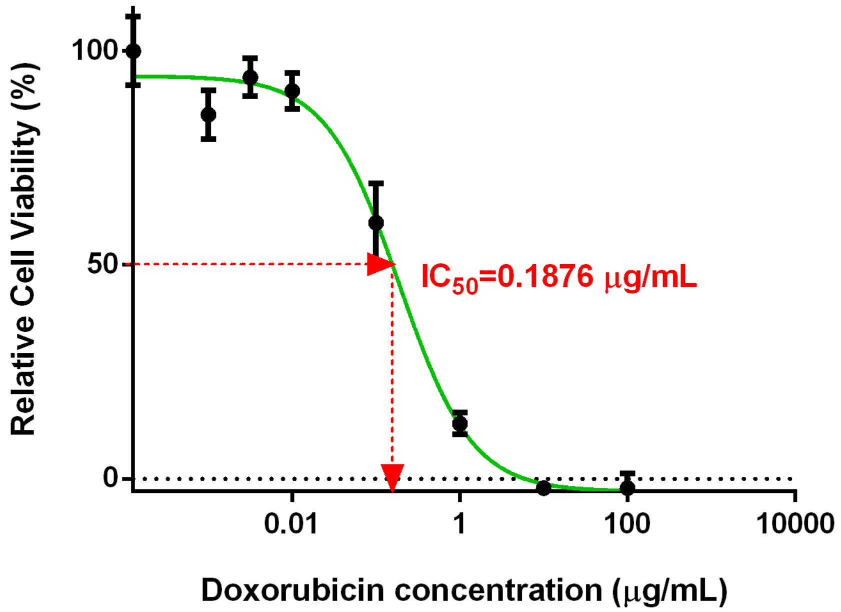


**Figure S2.** Dose response curve of Saos-2 cells grown in a tissue culture plate treated with Doxorubicin. (Cell seeding density = $2x10^4$ cells $well^{-1}$)
